# Acute and chronic gregarisation are associated with distinct DNA methylation fingerprints in desert locusts

**DOI:** 10.1101/018499

**Authors:** Eamonn B. Mallon, Harindra E. Amarasinghe, Swidbert R. Ott

**Affiliations:** Department of Genetics, University of Leicester, University Road, Leicester, LE1 7RH, United Kingdom.; Academic Unit of Cancer Genomics, Faculty of Medicine, University of Southampton, Southampton, SO17 1BJ, United Kingdom.; Department of Neuroscience, Psychology and Behaviour, University of Leicester, University Road, Leicester, LE1 7RH, United Kingdom.

**Keywords:** epigenetics, methylation-sensitive amplified fragment length polymorphisms, MS-AFLP, phase change, *Schistocerca gregaria*

## Abstract

Desert locusts (*Schistocerca gregaria*) show a dramatic form of socially induced phenotypic plasticity known as phase polyphenism. In the absence of conspecifics, locusts occur in a shy and cryptic solitarious phase. Crowding with conspecifics drives a behavioural transformation towards gregariousness that occurs within hours and is followed by changes in physiology, colouration and morphology, resulting in the full gregarious phase syndrome. We analysed methylation-sensitive amplified fragment length polymorphisms (MS-AFLP) to compare the effect of acute and chronic crowding on DNA methylation in the central nervous system. We find that crowd-reared and solitary-reared locusts show markedly different neural MS-AFLP fingerprints. However, crowding for a day resulted in neural MS-AFLP fingerprints that were clearly distinct from both crowd-reared and uncrowded solitary-reared locusts. Our results indicate that changes in DNA methylation associated with behavioural gregarisation proceed through intermediate states that are not simply partial realisations of the endpoint states.

## Introduction

Modification of neural DNA by cytosine methylation is emerging as an important mechanism in tailoring behavioural phenotypes to environmental conditions, including the social environment [1–4]. In most instances, however, the mechanistic role of DNA methylation in the chain of events from environmental signals to changes in behavioural phenotype is still poorly understood [4]. To what extent are changes in DNA methylation responsible for bringing about behavioural change, as opposed to serving to consolidate changes that first arose through other mechanisms? This is related to an even more fundamental question in understanding phenotypic transitions: are individuals in transition simply intermediates between more extreme endpoints, or are they better understood as ‘third states’? The answer may differ at different levels of analysis, as similar behavioural states may be underpinned by different mechanistic states.

Phenotypic plasticity is particularly common in insects, a fact implicated in their evolutionary success [5]. A striking example is provided by phase polyphenism in locusts. Locusts are grasshoppers (Acrididae) that can transform between two extreme phenotypes known as the *solitarious* and *gregarious phase*, which differ profoundly in morphology, physiology and behaviour [6]. Solitarious-phase locusts are cryptic and shy, and avoid conspecifics; gregarious-phase locusts are active and mobile and seek out conspecifics, causing them to aggregate in swarms. Several distantly related grasshopper species show phase polyphenism, with migratory locusts (*Locusta migratoria*) and desert locusts (*Schistocerca gregaria*) being amongst the most extreme and economically relevant. The sole direct environmental driver of phase change is the presence or absence of conspecifics. Solitarious desert locusts acquire gregarious behaviour within a few hours of forced crowding [7,8]. Behavioural solitarisation of long-term gregarious locusts is markedly slower, indicating a consolidation of the gregarious state with prolonged crowding. In desert locusts, phase state at hatching is additionally determined by trans-generational epigenetic inheritance [9].

Phase change in locusts provides an attractive model for addressing fundamental questions about the role of DNA methylation in behavioural plasticity. Neural DNA methylation could conceivably contribute to several different aspects of behavioural phase polyphenism. It could be part of the effector cascade that initiates behavioural change; it could contribute to the consolidation of gregarious behaviour that occurs with prolonged crowding within a lifetime; and it could contribute to the inheritance of phase state across generations.

The DNA in locust central nervous systems is heavily methylated by insect standards: in *S. gregaria,* methylation occurs on 1.6–1.9% of all genomic cytosines and on over 3% of the cytosines in exons [10,11]. These values are over tenfold higher than in honeybees, where methylation is implicated in caste polyphenism [12–14], suggesting that DNA methylation has important functions in locust behaviour. Two forms of (cytosine-5)-methyltransferases (DNMTs) are primarily responsible for DNA methylation in animals: DNMT3s, traditionally viewed as *de novo* methylation enzymes, and DNMT1s, which were thought to maintain the methylation state. The roles are not that clear cut, however [15]. In post-mitotic neurones, for example, DNMT1 and DNMT3 have overlapping roles in synaptic plasticity [16]. Unsuccessful attempts to identify *Dnmt3* transcripts in EST libraries from either *S. gregaria* or *L. migratoria* led to the suggestion that Acridids may have secondarily lost this gene, as have several other insect clades [10,17,18]. Wang and colleagues [19] reported the presence of a *Dnmt3* gene in the *L. migratoria* genome but did not include their evidence, so that further studies are needed to resolve whether a functional gene is present. In species that have lost DNMT3, *de novo* methylation might be accomplished by DNMT1 [18].

A practical difficulty is the lack of a sequenced reference genome for the desert locust and the huge genome sizes of Acrididae in general. In principle, several approaches are currently available for genome-wide studies of DNA methylation. Analysis of methylation-sensitive amplified fragment length polymorphisms (MS-AFLP), a gel electrophoresis-based technique, is still the most widely used approach in non-model organisms without a reference genome, despite its limitations [20]. MS-AFLP can only screen a limited number of loci simultaneously, and these remain anonymous, *i.e.,* differences in banding pattern cannot be directly linked to known genomic loci. In model species, MS-AFLP has been largely superseded by more powerful techniques that include whole-genome bisulfite sequencing (WGBS) and reduced representation bisulfite sequencing (RRBS). However, these techniques are not readily transferrable to organisms for which a reference genome is not yet available [21]. WGBS would presently be prohibitively difficult and expensive in desert locusts given the huge genome size of 8.55 Gb [22] and the expected abundance of repetitive elements (60% of the genome in *L. migratoria* [19]). RRBS is only sensible if the reads can be aligned against a reference genome, which presently precludes its use in desert locusts. The considerable technical complexities of RRBS [23] also make it less suitable for an initial screening of the time course of changes. In *L. migratoria,* a species with a fully sequenced genome of about 6.5 Gb, RRBS identified about 90 differentially methylated genes in the brains of solitarious and gregarious nymphs [19]; this relatively moderate number probably represents only the tip of an iceberg and highlights the challenges of RRBS in locusts even when both a reference genome and substantial laboratory resources are available. New NGS-based methods are now appearing that are designed specifically to overcome the limitations encountered in non-model species *(e.g.,* [21,24]). At present, the primary utility of MS-AFLP is to inform the design of more detailed follow-up experiments using such emerging NGS approaches.

For this study, we chose MS-AFLP to compare the neural DNA methylation fingerprints of desert locusts with identical parental histories, but different individual social rearing histories. The reasons for this choice were (i) the fact that MS-AFLP does not require a sequenced reference genome; and (ii) the relatively low cost of MS-AFLP, which enabled us to individually profile a reasonable number of animals for statistical analysis. The study was designed to answer three questions. First, do long-term solitarious and gregarious desert locusts show differences in their global pattern of neural DNA methylation, as was recently reported in migratory locusts [19]. This question is of interest because the two species are only distantly related and have evolved phase polyphenism independently [25]. The primary focus of our study, however, was on whether the neural DNA methylation fingerprint changes over a timescale of crowding that is sufficient for behavioural gregarisation. We therefore asked whether a day of crowding is sufficient to cause detectable changes in the neural DNA methylation fingerprint of solitary-reared locusts; and if so, whether the methylation fingerprint of these acutely gregarised locusts already resembles that of long-term gregarious locusts.

## Methods

### Locust rearing and treatments

Desert locusts (*Schistocerca gregaria* Forskål, 1775) were obtained from an inbred gregarious colony at Leicester. Solitarious-phase locusts were produced from this stock by transferring them within a day of hatching into individual cages and rearing them in visual, tactile and olfactory isolation [26]. All locusts were maintained on a diet of fresh seedling wheat and dry wheat germ under a 12:12 photoperiod.

All locusts were virgin adults sacrificed 17-21 days after the final moult. Long-term gregarious (LTG) locusts were removed from the colony as final larval instars, sexed, and set up as one all-male and one all-female cohort of 40 each in separate tanks (40 × 30 × 25 cm^3^) in the controlled-environment room that also housed the solitarious locusts. Solitarious locusts were offspring from a single gregarious mother (first-generation solitarious, 1GS). There were three treatment groups of four males and four females each: (i) *n* = 8 1GS locusts that never experienced crowding; (ii) *n* = 8 LTG locusts; and (iii) *n* = 8 behaviourally gregarised 1GS locusts. These were produced by placing four male and four female 1GS locusts in the tanks that housed the 40 LTG virgins of the respective sex for 24 h before sacrifice. Locusts were sacrificed by decapitation and immediate dissection under ice-cold saline. The brain (excluding the retinae) and the thoracic ganglia were dissected out and snap-frozen on dry ice.

### MS-AFLP analysis

Differences in DNA methylation patterns were detected by MS-AFLP analysis in *n* = 4 independent samples per treatment group, for a total of *N* = 12 samples. Each sample comprised the pooled brains and thoracic ganglia from one arbitrarily chosen male and female within the same treatment group. DNA was extracted with the QIAamp DNA Micro Kit (QIAGEN) following the manufacturer’s instructions.

#### Restriction digestion

The MS-AFLP protocol was based on [27]. For each sample of genomic DNA, one 500 ng aliquot was digested with EcoRI and MspI by combining 3 μl target DNA, 0.05 μl EcoRI (20 000 units/ml), 0. 25 μl MspI (20000units/ml), 1 μl 10 × NEBuffer 4 and 5.7 μl H_2_O); another 500 ng aliquot of genomic DNA was digested with EcoRI and HpaII by combining 3 μl target DNA, 0.5 μl EcoRI (20 000 units/ml), 0.5 μl HpaII (10 000units/ml), 1 μl 10 × NEBuffer 1 and 5.45 μl H_2_O) at 37°C for 3 h.

#### Adapter ligation

The EcoRI-MspI and EcoRI-HpaII restriction-digested products were ligated with EcoRI and HpaII-MspI adaptors (Table 1). The EcoRI adaptor was prepared from 5 μl EcoRI-F and 5 μl EcoRI-R, mixed in a final concentration of 5pmol μl^−1^ each; the HpaII-MspI adaptor was prepared from 25 μl HpaII-MspI-F and 25 μl HpaII-MspI-R, mixed in a final concentration of 50pmol μl^−1^ each. Both mixes were incubated at 65°C for 10 min. For ligation, 3 μl digested product was combined with 7 μl of ligation reaction mixture (1 μl EcoRI adapter, 1 μl HpaII-MspI adapter, 0.25 μl T4 DNA ligase (400 000units/ml), 1 μl 10 × T4 ligase buffer (New England Biolabs) and 3.75 μl H_2_O) at 37°C for 3 h and then left overnight at room temperature. The ligation products were diluted with 100 μl of H_2_O and used as the template for pre-amplification.

**Table 1.**
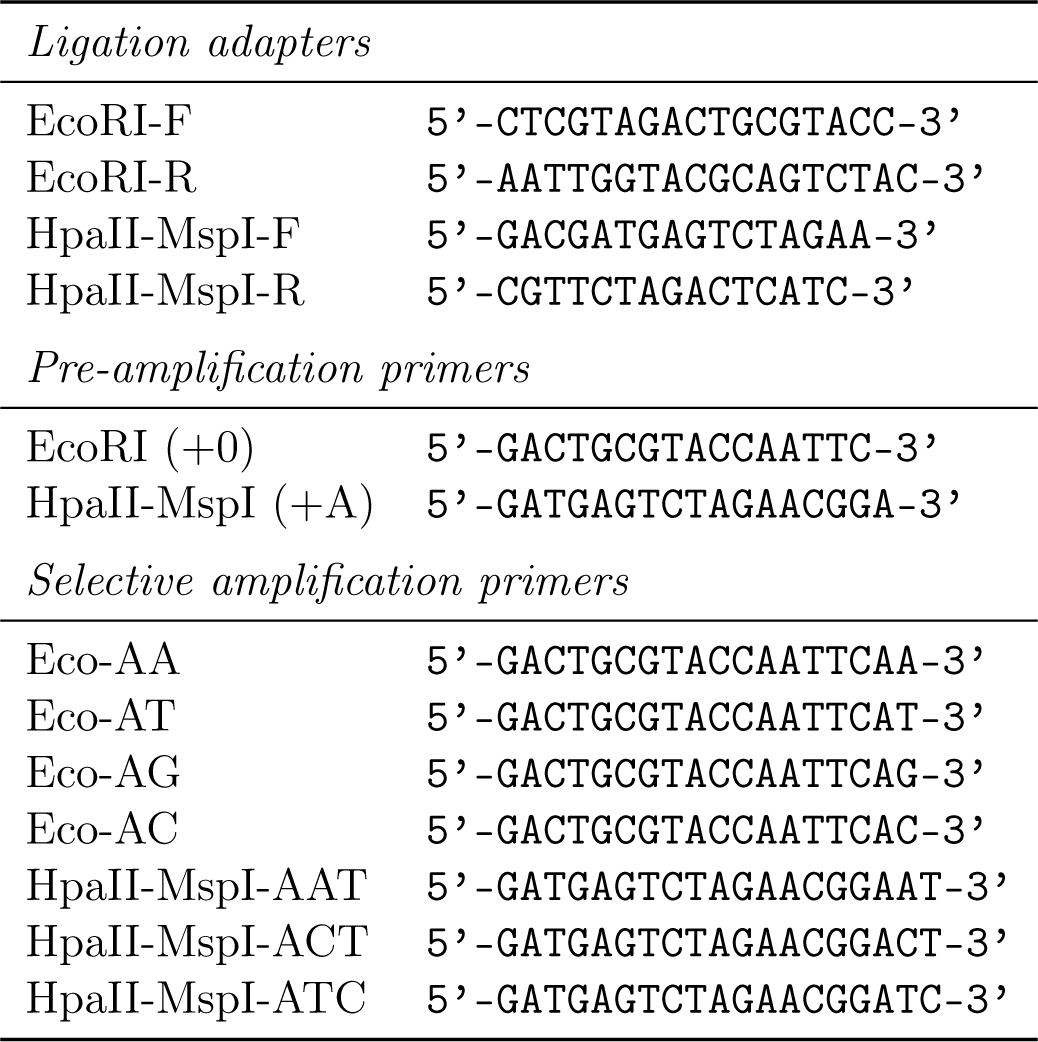
Sequences of ligation adapters, pre-amplification primers and selective amplification primers.

#### Pre-amplification

The pre-amplification PCR used 1 μl of ligation product with 1 μl each of EcoRIpre and HpaII-MspIpre primers (10pmol ml^–1^; Table 1), and 7 μl of the reaction mix (0.8 μl 2.5 mM deoxynucleotide triphosphates (dNTPs), 1 μl 10 × Paq5000 Hot Start Reaction Buffer, 0.3 μl Paq5000 Hot Start DNA Polymerase (500 units), 0.8μl 25 mM MgCl_2_, 4.1 μl sterile H_2_O). The PCR conditions were 94°C for 2 min, followed by 20 cycles of 94°C for 30 s, 60°C for 1 min and 72°C for 1min, followed by a final extension of 5 min at 72°C. 3 μl of each PCR product was run on 3% agarose gel and appearance of a smear of DNA on the gel indicated that the preamplification PCR was successful.

#### Selective amplification

Seven μl of PCR products were diluted with 93 μl of H_2_O and used as the template for selective amplification. We used four different selective EcoRI primers and three different HpaII-MspI primers (Table 1), giving twelve unique EcoRI/HpaII-MspI primer pair combinations. In order to reduce the number of bands in the subsequent gel electrophoresis to a manageable number, each primer combination was used in a separate PCR, giving twelve PCR products per sample that were subsequently run on separate gels. The selective PCR reaction mixtures contained 1 μl preamplified product, 1 μl each of one of the HpaII-MspI primers and of one of the EcoRI primers (10 pmol ml^−1^) and 7 μl reaction mix (same as used for pre-amplification). PCR conditions were: (i) 94°C for 2 min; (ii) 13 cycles of 30 s at 94°C, 30 s at 65°C (0.7°C reduction per cycle) and 1 min at 72°C; (iii) 23 cycles of 30 s at 94°C, 30 s at 56°C and 1 min at 72°C; and (iv) a final extension at 72°C for 5 min followed by a holding step at 4°C.

#### Gel electrophoresis

PCR products were diluted with 100 μl H_2_O. 10 μl of diluted PCR product was mixed with 3 μl 1x loading buffer (Elchrom Scientific, Cham, Switzerland) and run on 9% poly(NAT) gels (Elchrom) on an Elchrom Origins electrophoresis system (120V, 81 min at 55°C). Gels were stained in the dark with SYBR^®^ Gold (Invitrogen; 1:10 000 in TAE buffer) followed by destaining in 100 ml TAE buffer alone.

#### Statistical analysis

Bands were scored automatically as either present or absent by the Java program GelJ [28]. Positions of loci are reported as molecular weights (base pairs). GelJ matches bands across different gel lanes based on a user-specified tolerance value (in base pairs) below which bands are considered identical. To ensure that our results were not sensitive to this arbitrary tolerance level, we generated matrices of band scores using tolerance values of 1-40 using custom R scripts. The resulting matrices were analysed for differentiation between groups by principal coordinates analysis (PCoA) and by analysis of molecular variance (AMOVA) in the R package *msap* [29]. We investigated the sensitivity of the Φ_*ST*_ value to the choice of tolerance value. This identified a broad range of tolerances which gave the same robust result (see Results and Discussion). Finally, to ensure that the Φ_*ST*_ values generated across this range of tolerance values were not due to chance, we compared Φ_*ST*_ from the real data with bootstrapped Φ_*ST*_ values from data generated by random sampling with replacement (*N* = 1000).

All R scripts used are available at https://dx.doi.org/10.6084/m9.figshare.3168760.v1. All statistical analysis was carried out in R 3.2.3 [30].

## Results and Discussion

Our analysis of the MS-AFLP patterns across solitary-reared, 24 h crowded and crowd-reared locusts indicated clear differences between the three groups. This result was robust across a range of band scoring tolerances — the distance up to which bands in a given gel position are matched as identical between samples (Figure 1). As expected, tolerance had some effect on our results. At low tolerances (<10bp), a large fraction of the bands within each sample are treated as unique, leading to low calculated levels of differentiation between the groups (Φ_*ST*_). Tolerances in the range of 10-25 bp produced robust Φ_*ST*_ values of approximately 0.2. The principal coordinate analysis for each of these tolerance levels is shown in Supplementary Figures S1-S3. Above this range, Φ_*ST*_ began to drop. This is to be expected, as bands that represent genuinely different fragments will now be grouped together, leading to less apparent differentiation between groups. This pattern of increasing and then decreasing Φ_*ST*_ is not due to chance as the bootstrapped data, generated by random sampling with replacement, do not show a similar pattern. Importantly, across the entire range of tolerances, the Φ_*ST*_ values obtained in the real data (red points in Figure 1) are well outside the bootstrapped Φ_*ST*_ distributions (grey points and black boxplots in Figure 1).

**Figure 1.**
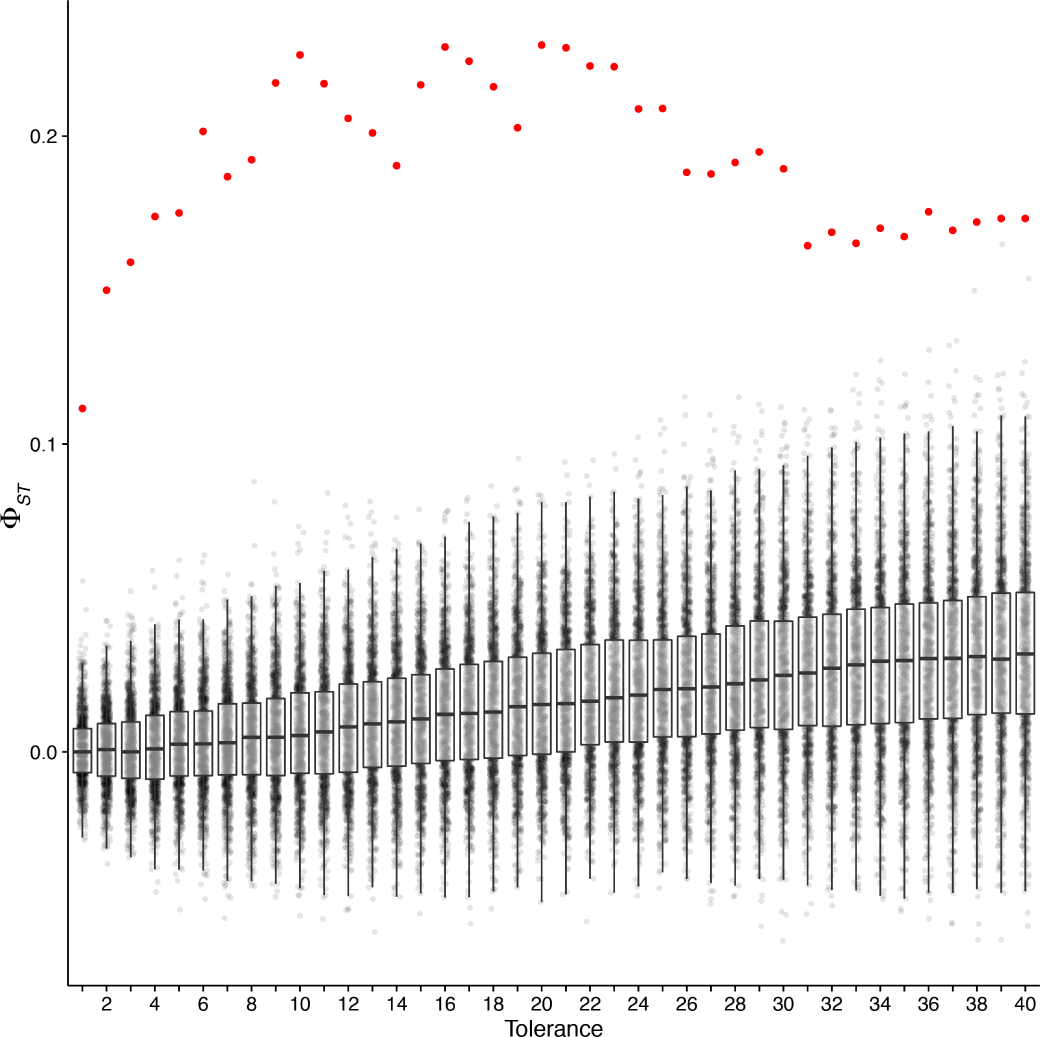
Sensitivity of our analysis to the tolerance value used for matching bands between samples. Φ_*ST*_ represents the apparent degree of differentiation between the groups (solitary-reared, crowd-reared, and solitary-reared crowded for 24 h) and is plotted over the range of tolerance values (in base pairs). The red points represent Φ_*ST*_ values calculated from the real data, the grey points those calculated from *N* = 1000 bootstrapped data sets (generated by random sampling with replacement).

The following analysis is based on a tolerance value of 10 bp. This identified 294 unique AFLP bands (loci); of these, 282 were identified as methylation-susceptible based on different digestion patterns with HpaII and MspI, and 162 showed different banding patterns between individual samples (MS-polymorphic loci). Solitary-reared locusts had a slightly lower proportion of unmethylated loci than crowd-reared locusts (11.8% vs. 16.3%) and a slightly higher proportion of hypermethylated loci (65.6% vs. 55.9%). Acutely crowded solitary-reared locusts showed proportions of methylation that were intermediate (Table 2). The three treatment groups showed significant multi-locus differentiation in their methylation fingerprints (AMOVA, Φ_*ST*_ = 0.2264, *p* = 0.0002). Figure 2 gives a simplified representation of the multi-locus differentiation between the samples. The two axes represent the first two principal coordinates, which together explained 35.1% of the total variation.

**Figure 2.**
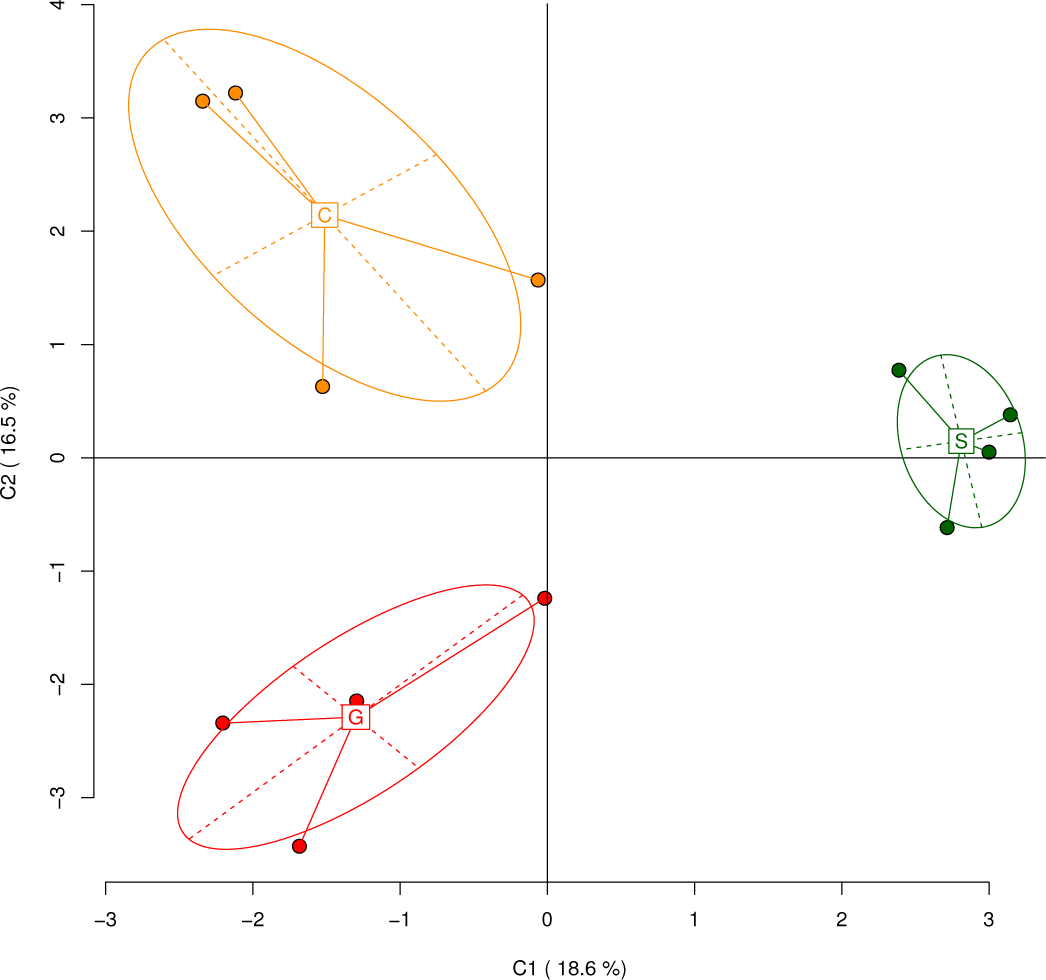
Principal Coordinate Analysis (PCoA) of epigenetic differentiation between uncrowded solitary-reared locusts (S), long-term gregarious locusts (G) and solitary-reared locusts crowded for 24 h (C), as identified by MS-AFLP (10 bp band matching tolerance). The first two coordinates (C1, C2) are shown with the percentage of variance explained by them. Group labels show the centroid for each group, points correspond to individual MS-AFLP samples, ellipses represent their average dispersion around the group centroids.

**Table 2.** Proportion of methylation-sensitive restriction band patterns found in the CNS of locusts of different phase state, and their corresponding methylation status; methylated cytosines are indicated in bold type. ^a^ + and — indicate the presence and absence, respectively, of a band following digestion with HpaII or MspI. ^b^ may indicate methylation of either outer or both cytosines on one strand. ^c^ HPA— / MSP— was taken to indicate hypermethylation rather than absence of target due to a genetic mutation [29].

| Banding pattern <sup>a</sup> | Methylation |  | solitary | 24 h crowded | crowd-reared |
| --- | --- | --- | --- | --- | --- |
| HPA+ / MSP+ | none: | 5′-CCGG<br>GGCC-5′ | 11.8% | 15.0% | 16.3% |
| HPA+ / MSP– | hemi: <sup>b</sup> | 5′-CCGG<br>GGCC-5′ | 10.1% | 11.6% | 13.4% |
| HPA– / MSP+ | full internal: | 5′-CCGG<br>GGCC-5′ | 12.5% | 12.2% | 14.4% |
| HPA– / MSP– | hyper: <sup>c</sup> | 5′-CCGG<br>GGCC-5′ | 65.6% | 61.2% | 55.9% |

A pair-wise comparison between crowd-reared and solitary-reared locusts identified significant epigenetic differentiation (Φ_*ST*_ = 0.2810, *p* = 0.0291), indicating that phase change in desert locusts entails modification of the neural DNA methylation pattern. This is maybe the least surprising of our results, considering that differences in brain DNA methylation between longterm phases have been previously reported in the distantly related migratory locust *L. migratoria* [19]. The differences observed in our experiment arose within a single generation, because we used solitary-reared locusts that were the direct offspring of long-term gregarious parents. It would now be interesting to see whether isolation over multiple generations further deepens the epigenetic differences between the two phases.

Our key finding, however, is that crowding solitary-reared locusts for 24 h resulted in a neural DNA methylation fingerprint that was distinctly different both from uncrowded solitary-reared locusts Φ_*ST*_ = 0.2381, *p* = 0.0283) and from crowd-reared locusts (Φ_*ST*_ = 0.166, *p* = 0.0288). This uncovers a disjunct between the global neural DNA methylation pattern and the behavioural phase state. Although one day of crowding is sufficient to establish gregarious behaviour [7,8], we find that the neural MS-AFLP fingerprint is at this point still markedly different from that in long-term gregarious locusts. The time-course of behavioural gregarisation has been characterised primarily through work carried out in juveniles, where gregarious behaviour is established within 4h of crowding *(e.g.,* [7,31]). Therefore, the difference that we observed between the methylation fingerprints of 24 h crowded and crowd-reared adult locusts could reflect an incomplete or slower behavioural gregarisation response in adults, *e.g.,* due to age-related limits of plasticity. Bouaïchi and colleagues [8] reported that very young adults, 2 days post-fledging, responded more strongly to crowding than older adults, 7-35 days post-fledging (‘Experiment 1’ in [8]). However, this effect was not robust (not evident in ‘Experiment 2’ of [8]), and after 8h of crowding, adult locusts were found behaviourally indistinguishable from crowd-reared locusts. To the extent that they have been documented, age-related differences in the gregarisation response are minor and set against a clear behavioural transformation in less than 24 h, irrespective of age. We therefore expect that the result of this study generalises to juveniles.

Interestingly, the data points from 24 h crowded samples were set apart from the solitary-reared samples and the crowd-reared samples by shifts along both of the first two principal coordinate axes (Figure 2). Although the exact position of the group centroids relative to the two axes depended on the tolerance value used in the band scoring algorithm, a clear triangular separation was maintained across the entire range of sensible tolerance values (Supplementary Figures S1-S3). In other words, the three groups never fell along a single line in the PCoA plots. We interpret this as evidence that the methylation patterns seen after 24 h of crowding are not simply intermediate between the two extremes, but reflect a distinct transitional epigenetic state.

The further changes in methylation that occur only some time after the first 24 h of crowding must then be mechanistically unrelated to the transition to, or expression of, gregarious behaviour. Previous behavioual studies have shown that the resilience of gregarious behaviour to re-isolation increases with time spent in crowded conditions [26,32]. When solitarious locusts are re-isolated after 24-48 h of crowding, they return to fully solitarious behaviour within 8 h. Long-term gregarious locusts, however, solitarise only partially when isolated for four days as final instar nymphs. Some of these late changes in methylation pattern may therefore represent consolidation mechanisms by which neurochemically mediated rapid changes in behaviour [31] become more stable with time. However, differential DNA methylation may also underpin long-term phase differences in the CNS that are not directly responsible for generating phase-specific behaviour but represent adaptations to the respective life styles.

Our present results add to previous evidence for the existence of mechanistically distinct transitional phase states at different levels from the molecular to the behavioural. On a neurotransmitter level, serotonin concentrations in the CNS show a marked transient increase in the thoracic ganglia within the first few hours of crowding that has been causally linked to the transition to gregarious behaviour [31]. This is followed by an equally transient increase in the brain around 24 h [33]. On a neuronal level, acute and chronic crowding also have differential effects on serotonin-sythesising neurones, with one set of neurones responding to acute crowding with increased serotonin expression and a distinct set showing decreased serotonin expression in the long term [34]. Even on a behavioural level, when forming new associations between unfamiliar odours and toxic food, recently gregarised locusts differ from both long-term phases, in a way that matches their respective distinct ecological requirements [35–37].

In conclusion, our results demonstrate that phase change in desert locusts is associated with distinct short-and longterm shifts in the neural DNA methylation fingerprint. A purely associative study like ours cannot prove causal connections between methylation and behaviour. However, an important consequence of our findings for future studies is that uncovering such causal connections will require analyses of transitional stages rather than only comparisons of endpoints [19] because the transitional states are not simply partial realisations of the endpoints. Newly emerging NGS-based methods that are designed specifically to enable epigenome-wide association studies in non-model species *(e.g.,* [21,24]) hold great promise for following up the present results to identify specific changes at the level of identified methylation sites.

## Acknowledgments

Supported by research grants F/09 364/K from the Leverhulme Trust, UK (to SRO), BB/L02389X/1 from the Biotechnology and Biological Sciences Research Council, UK (to SRO) and NE/N010019/1 from the Natural Environment Research Council, UK (to EBM). We thank Alice Round and Carl Breaker for managing our locust facilities.

## Additional Information

### Authors’ contributions

SRO and EBM conceived the study and designed the experiments. SRO carried out the animal treatments and dissections. HEA carried out all MS-AFLP bench-work and initial gel analysis and prepared the first draft. EBM and SRO performed statistical analyses. SRO wrote the final draft with input from HEA and EBM. All authors gave final approval for publication.

### Competing financial interests

The authors declare no competing financial interests.

